# A structure-based modelling approach identifies effective drug combinations for RAS-mutant acute myeloid leukemia

**DOI:** 10.1101/2025.04.29.651188

**Authors:** Luke Jones, Oleksii Rukhlenko, Tânia Dias, Hiroaki Imoto, Ciardha Carmody, Kieran Wynne, Boris N. Kholodenko, Jonathan Bond

## Abstract

Mutations activating RAS/RAF/MEK/ERK signaling are associated with poor outcome in acute myeloid leukemia (AML), but therapeutic targeting of this pathway is challenging. Here, we employ a structure-based, dynamic RAS pathway model to successfully predict RAF inhibitor (RAFi) combinations which synergistically suppress ERK signaling in *RAS*-mutant AML.

Our *in silico* models predicted therapeutic synergy of two iterations of conformation-specific RAF inhibitors: Type I½ + Type II and Type I + Type II. Predictions were validated *in vitro* in AML cell lines and patient samples, with synergy verified by the Loewe Additivity model. Lifirafenib (Type II) + encorafenib (Type I½) was highly synergistic against both *NRAS*– and *KRAS*-mutant lines, while synergy of lifirafenib + SB590885 (Type I) was specific to *NRAS*-mutants. Immunoblotting confirmed that combination efficacy correlated strongly with decreased RAS pathway activation.

Leveraging the pharmacokinetic predictions of our *in silico* model, both combinations were then assessed in a pre-clinical *NRAS*-mutant AML patient-derived xenograft (PDX) model, showing significantly improved leukaemia growth delay and event-free survival compared with single agent approaches. Assessment of leukemia burden in bone marrow and spleen during treatment further showed site-specific efficacy against circulating and spleen-resident blasts for both combinations.

In summary, we report that our structure based-modelling approach can effectively identify novel, non-obvious, and well-tolerated RAFi combinations that are highly effective against *in vitro* and *in vivo* models, thereby suggesting alternative potential therapeutic strategies for high-risk *RAS*-mutant AML.

## Introduction

Mutations that activate RAS signaling are common in myeloid malignancies and are typically associated with a proliferative phenotype and aggressive disease.^1–4^ These mutations are frequent in acute myeloid leukemia (AML) regardless of age, with evolving evidence suggesting that RAS pathway activation alters therapy responsiveness. For example, *RAS* mutations are associated with worse outcome following hypomethylating agent (HMA) treatment, whether AZA alone or combined Ven-AZA.^5–9^ Additionally, *RAS* mutations have also been implicated in resistance to targeted therapies, including FLT3 and IDH1/2 inhibitors.^10–12^

Further, recent reports have linked *KRAS* mutations with inferior survival within *KMT2A*-rearranged AML cohorts.^13,14^ Mutations activating RAS signaling were also recently shown to be the most common co-operative mutations in *FUS::ERG* AML, which is associated with poor outcome.^15^ Additionally, mutations in regulators of RAS signaling, *NF1* and *PTPN11*, have both been reported as independent predictors of poor prognosis. Activating mutations in *RAS* have also been associated with poor outcome and chemoresistance in other hematological malignancies, including T-acute lymphoblastic leukemia (T-ALL).^16^

These prognostic associations have driven efforts to address the currently unmet need of targeting aberrant RAS signaling in AML and other cancers. Prior to the recent development of direct inhibitors of RAS, which are largely untested in the context of myeloid malignancies, RAS proteins were considered ‘undruggable’. Therefore, efforts to inhibit aberrant RAS signaling have focused on targeting factors upstream and downstream of RAS. Of these, MEK inhibition has shown most promise, exhibiting modest efficacy in preclinical models by reducing downstream ERK phosphorylation and proliferation.^17,18^ Targeting RAF, the kinase immediately downstream of RAS, has been less well investigated due to the phenomenon of paradoxical activation, whereby single-agent RAF inhibitors cause increased pathway activity.^19–21^

We recently reported a structure-based *in silico* modeling approach for identifying non-obvious RAF kinase inhibitor combinations that effectively suppress oncogenic RAS signaling.^22^ As with other kinases, RAF toggles between inactive and active conformations which differ by the relative positions of the highly conserved DFG motif and αC helix. ATP-competitive RAF inhibitors are classified based on their preference for binding to alternate (IN or OUT) conformations of the RAF kinase ATP-binding pocket (IN and OUT positions correspond to active and inactive kinase conformations, respectively) (Figure 1).^23–25^ There are three distinct RAF inhibitor types, specifically targeting: αC-IN/DFG-IN (denoted CI/DI, Type I), αC-OUT/DFG-IN (CO/DI, Type I½,), and αC-IN/DFG-OUT (CI/DO, Type II).

**Figure 1.**
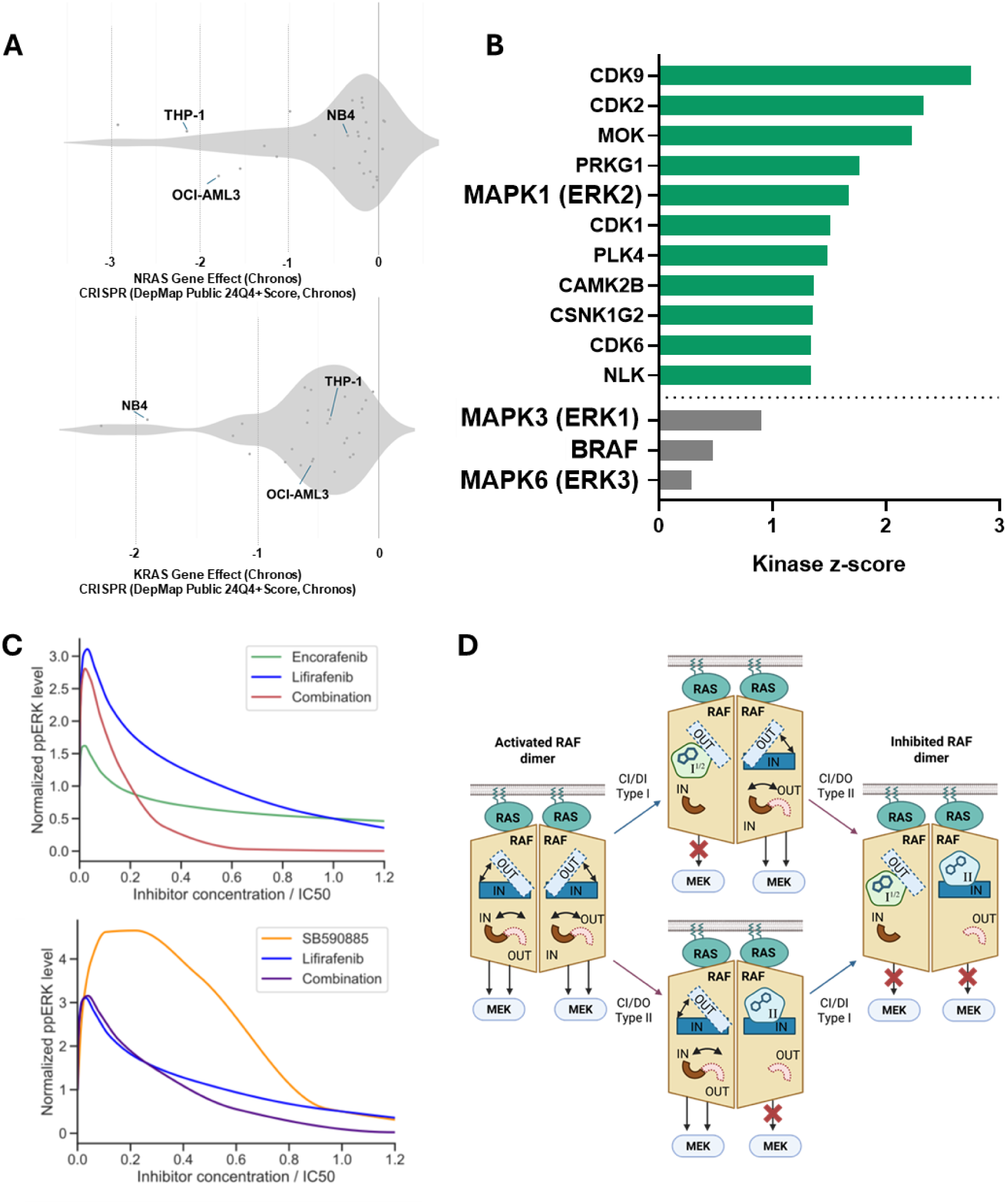
Structure-based modeling predicts RAF inhibitor combinations effective against *RAS*-mutant AML. **(A)** Gene essentiality scores for *NRAS* (top panel) and *KRAS* (bottom panel) across AML cell lines, ranked using data available from CRISPR screens in DepMap (https://depmap.org/portal/). **(B)** Kinase Set Enrichment Analysis (KSEA) comparing kinase enrichment in *RAS*-mutant (n=5) versus *RAS*-WT (n=3) cell lines. Green bars indicate kinases significantly enriched in *RAS*-mutant cells. Grey bars represent RAS pathway components enriched in *RAS*-mutant cells that did not meet the statistical threshold (P < 0.1). **(C)** Steady-state phosphorylated ERK (ppERK) responses to RAF inhibitor treatments. Inhibitor concentrations were normalized to their respective IC₅₀ values. The equilibrium dissociation constant (Kd) for SB590885 (type I) is reported as 0.3 nM. For encorafenib (type I½) and lifirafenib (type II), Kd values are not publicly available; therefore, we approximated Kd using available EC₅₀ values from SelleckChem. The Kd values used in the model were 0.3 nM for SB590885, 4 nM for encorafenib, and 23 nM for lifirafenib. **(D)** Cartoon depicting the binding preferences and effects on downstream signaling upon treatment with Type I½, Type II and Type I½ + Type II RAF inhibitors.

Importantly, we and others showed that the paradoxical activation of downstream ERK signaling in response to single-agent RAF inhibitors is primarily due to RAF dimer asymmetry. As each protomer exists in an alternate conformation (IN-OUT), use of a single RAF inhibitor will bind one protomer and cause allosteric activation of the other.^26–29^ Our structure-based modelling approach showed that RAF dimer asymmetry is a targetable vulnerability, with the combination of structurally different RAF inhibitors abrogating paradoxical activation and significantly reducing downstream pathway activation and cell viability in *RAS*– and *RAF*-mutant solid tumor models.^22,30^

Given the prevalence and poor prognosis linked to RAS signaling mutations in AML, we now apply our modelling approach to leukemia to address an unmet clinical need. By refining and adapting our structure-based approach by integrating AML-specific kinase pathway activities, we have now identified novel combination approaches that effectively kill *RAS*-mutant AML *in vitro* and in preclinical *in vivo* models.

## Methods

### Cell lines

Cell lines were acquired from American Type Culture Collection (ATCC®) or Deutsche Sammlung von Mikroorganismen und Zellkulturen (DSMZ). HL-60, OCI-AML3, THP1 and NB4 cells were cultured in RPMI 1640 supplemented with GlutaMAX (2 mM) and 10% fetal bovine serum. ML-2 cells were cultured RPMI 1640 with GlutaMAX (2 mM) and 20% fetal bovine serum. All cell lines were cultured at 37°C and 5% CO_2_. The identities of all cell lines were validated using the cell line authentication service provided by Eurofins genomics (https://eurofinsgenomics.eu/en/genotyping-gene-expression/applied-genomics-services/cell-line-authentication/), using the Applied BiosystemsTM AmpFLSTRTM IdentifilerTM Plus PCR Amplification Kit system. Cell lines in culture were tested regularly (at least every 3 months) for mycoplasma using Lonza’s MycoAlert® Mycoplasma Detection Kit (LT07-710) according to the manufacturer’s instructions.

### Animal models

NSGS (NOD-scid IL2Rgnull-3/GM/SF) breeding pairs were purchased from Charles River Laboratories. Breeding was performed in-house in the Biomedical Facility (BMF) at University College Dublin. All mice were maintained under SPF conditions within the BMF. Male and female progeny were used in equal number for all *in vivo* experiments. All animal experiments were approved by the UCD Animal Research Ethics Committee (Approval AREC-21-04-Bond) and the Health Products Regulatory Authority (HPRA; Authorisation AE18982/P199).

### Inhibitors

GDC-0879 (#S1104), vemurafenib (PLX4032, #S1267), dabrafenib (#S2807), PLX7904, sorafenib (#S7397), TAK-632 (#S7291), tovorafenib (TAK-580, #S7121), RAF709 (#S8690) and RAF265 (CHIR-265, #S2161) were purchased from Selleck Chemicals LLC, Houston, TX, USA. Lifirafenib (BGB-283, # HY-18957), encorafenib (LGX818, HY-15605) and SB590885 (#HY-10966) were purchased from MedChemExpress EU, Sollentuna, Sweden. For *in vitro* use, all inhibitors were dissolved in DMSO (10mM) and stored at –80°C. For *in vivo* use, lifirafenib and encorafenib were suspended in 0.5% carboxymethylcellulose with 0.5% Tween-80 in sterile water and dissolved via sonication. SB590885 was dissolved with stepwise addition of 2% *N*,*N*-dimethylacetamide and 2% Cremophor EL in acidified water (pH = 5.0).

### Proteomics analysis

Mass spectrometry (MS) output data was searched using MaxQuant (release 2.0.1.0) against the *Homo sapiens* subset of the Uniprot Swissprot database with specific parameters for TIMS data dependent acquisition (TIMS-DDA). The MaxQuant output file was imported into the Perseus (version 1.6.15.0) environment for protein quantification. Phosphoproteome data was similarly exported and transformed to enable analysis of phosphorylation level and residue position. Full details of mass spectrometry analysis are provided in Supplementary Methods.

### Structure-based modelling of RAS/RAF/MEK/ERK signaling

To predict ERK phosphorylation dynamics in the venous blood, spleen, and bone marrow compartments, we integrated a physiologically based pharmacokinetic (PBPK) model (see Supplementary Methods) with our structure-based MAPK signaling model.^31^

In the previous structure-based models^22,29,31,32^, RAF inhibitor concentrations were fixed over time. Here, we dynamically updated inhibitor concentrations by the time-resolved drug concentration profiles predicted by the PBPK model for the 14 relevant compartments (see Supplementary Methods). This allowed the integrated model to capture the temporal dynamics of the drug exposure.

To match the experimental dosing regimen in mice, the model accounted for (i) SB590885 (Type I RAFi) administered intraperitoneally every 24 hours, (ii) encorafenib (Type I½ RAFi) administered orally every 24 hours, and (iii) lifirafenib (Type II RAFi) administered orally every 12 hours, both as single drugs and in combinations.

The integrated model was built using the rule-based PySB framework.^33^ The SBML files of the model used in this study are available in Supplementary Information.

### Assessment of *in vivo* efficacy

The AML005a PDX was obtained from the Baylor College of Medicine Patient-Derived Xenograft Core. AML005a cells (1×10^6^) were inoculated intravenously (i.v.) into unconditioned NSGS mice (6-8 weeks of age). Mice were bled (via tail vein) weekly to determine the percentage of human CD45^+^ (hCD45^+^) cells vs. mouse CD45^+^ (mCD45^+^) cells in peripheral blood (PB) using flow cytometry. When the median engraftment reached ≥1% hCD45^+^ cells, mice were assigned to treatment groups (n=8). Mice were treated with either lifirafenib (10 mg/kg PO twice daily), encorafenib (20 mg/kg PO once daily), SB590885 (25 mg/kg IP once daily), combined lifirafenib + encorafenib, combined lifirafenib + SB590885 or vehicle control. Representative mice were euthanized at day 10, with BM and spleen harvested to assess leukaemia burden and on-target efficacy of treatments. Mice were maintained in the experiment until they reached event, deemed 25% hCD45^+^ cells in PB.

Event-free survival (EFS) for individual mice was calculated as the number of days from treatment initiation until event, with the time of event calculated by interpolating between bleeds directly pre– and post-event, assuming log-linear growth. Leukemia growth delay (LGD) was measured as treatment minus control (T-C) in days. To evaluate interactions between drugs *in vivo*, therapeutic enhancement was considered if the EFS of mice treated with the combination treatment was significantly greater (P < 0.05) than those induced by both single agents.

Methods relating to immunoblotting, proteomics, Kinase Set Enrichment Analysis (KSEA), cell viability measurement, *RAS* mutation analysis and computational modeling are included in the Supplemental Methods section.

## Results

### Structure-based modelling identifies dual RAFi combinations for RAS-mutant AML

We first evaluated the contribution of oncogenic *RAS* mutations to the proliferation and survival of leukemia cells. We assessed *RAS* essentiality in AML cell lines using DepMap^34^ (https://depmap.org/portal) to help us select the optimum *in vitro* models. Of AML cell lines with available CRISPR-based dependency screens, THP-1 and OCI-AML3 ranked second and third respectively in *NRAS* dependence (Figure 1A). Assessment of *KRAS* dependency ranked NB4 as the second-most *KRAS*-dependent AML cell line (Figure 1A). To these, we added two additional cell lines, HL-60 (*NRAS*-mutant) and ML-2 (*KRAS*-mutant). Full details of cell lines used in this study are provided in Supplementary Figure S1A.

We next generated basal phospho-proteomic profiles for all five *RAS*-mutant cell lines. These data were first used to confirm RAS hyperactivation by using kinase set enrichment analysis (KSEA) to compare *RAS*-mutant cell lines with *RAS*^WT^ cell lines (MOLM-13, MV4;11 and PL-21). In line with RAS hyperactivation, this analysis showed significant enrichment of MAPK1 (ERK2) activity (Figure 1B). Further, other RAS/RAF/MEK/ERK pathway members (MAPK3/ERK1, BRAF and MAPK6/ERK3) also showed modest albeit non-statistically significant enrichment in *RAS*-mutant cells (Figure 1B).

Based on the strong dependence of these AML cell lines on oncogenic RAS signaling, we set out to apply our structure-based model of RAS/RAF/MEK/ERK signaling, integrated with a PBPK model of drug kinetics in mouse tissues, to RAS-mutant AML. This integrated model captures key intracellular processes, including ERBB receptor dimerization, recruitment of adaptor proteins to the plasma membrane, RAS activation through SOS1-mediated GDP-to-GTP exchange, RAF dimerization, MEK and ERK activation, and negative feedback loops from phosphorylated ERK to RAF, SOS1, and ERBB receptors. It also accounts for the kinetics of RAF inhibitors in mouse tissues and their allosteric interactions with various BRAF and CRAF protein complexes. The allosteric effects of conformation-specific drugs are quantitatively modeled using thermodynamic factors.^29^

The integrated model predicts the combinations of conformation-specific RAF inhibitors to be effective against *RAS*-mutant AML by durably and substantially suppressing ERK signaling. The predictions exploit the asymmetry of RAF dimers, by targeting the most thermodynamically favorable conformation of each monomer before and after inhibitor binding. In line with previous data observed for *RAS*– and *RAF-*mutant solid tumors, modelling predicted two combination types as potentially effective in a leukemic context: Type I + Type II and Type I½ + Type II (Figure 1C and D, Supplementary Figure S1B).

### Combining Type I½ /Type II RAF inhibitors is synergistic against NRAS and KRAS mutant AML in vitro

We first assessed the single-agent *in vitro* efficacy of RAF inhibitors representing both Type I½ (CO/DI, n=5) and Type II (CI/DO, n=3) inhibitor classes (Supplementary Figure S2A). Inhibitors with the lowest average IC_50_ across all *RAS*-mutant AML cell lines were chosen as candidates for combination testing. Lifirafenib (BGB-283), TAK-632 and sorafenib were chosen as candidate Type II inhibitors, while encorafenib and vemurafenib were taken forward as representative Type I½ inhibitors. While results were largely similar across the tested variations of Type I½ / Type II combinations (Supplementary Figure S2B), lifirafenib (Type II) and encorafenib (Type I½) was chosen for further experiments based on both observed synergy and clinical relevance, with encorafenib being FDA-approved and lifirafenib currently being evaluated in Phase II trials.

Both fixed-ratio and matrix (6×6) combinations were performed for all cell lines and synergy assessed using the Loewe Additivity method (>10 = synergy, 10 to –10 = additivity, <-10 = antagonism), chosen as both inhibitors share the same molecular target. In line with our structural biological predictions, this combination significantly reduced cell viability compared with single agents in all cell lines tested (Figure 2A). Strong synergy was observed in all *NRAS*-mutant cell lines, with Loewe Additivity scores of 31.1, 26.2 and 22.8 for HL-60, OCI-AML3 and THP-1, respectively. Synergy was also observed in the *KRAS*-mutant NB4 cell line (score = 18.7), while the combination was additive in the *KRAS*-mutant ML-2 cell line (score = 7.6) (Figure 2B).

**Figure 2.**
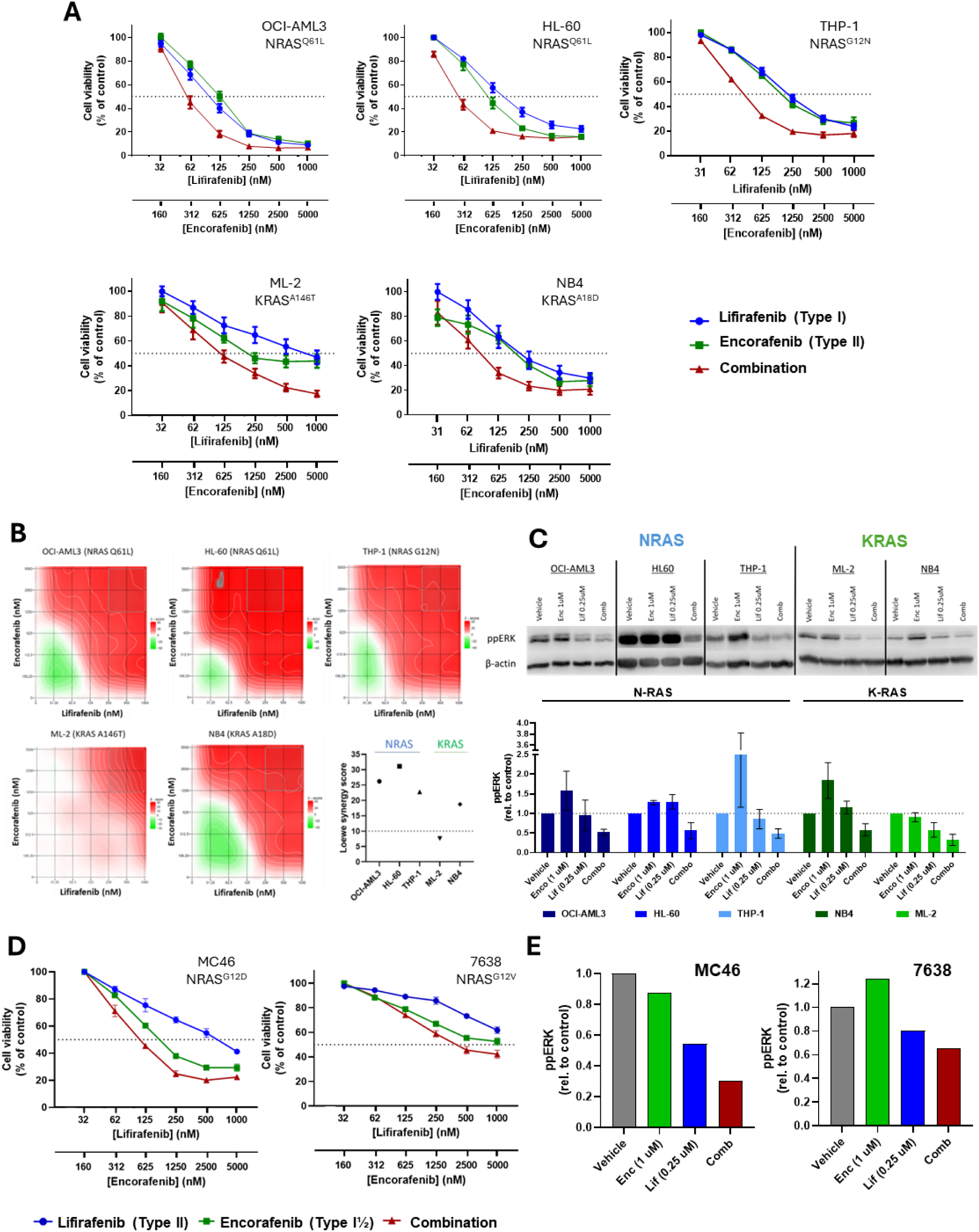
Combining Type I½ + Type II RAF inhibitors is synergistic against *RAS*-mutant AML cell lines and patient samples *in vitro*. **(A)** Cell viability in response to lifirafenib (Type II) + encorafenib (Type I½) compared with each single agent against five *RAS-*mutant AML cell lines. Cell viability was determined using AlamarBlue at 72 hours post-treatment. Data points represent mean ± SEM from 3 replicate experiments. **(B)** Synergy plots for lifirafenib + encorafenib in all five cell lines tested. Synergy was determined using the Loewe Additivity method, calculated using the SynergyFinder web application (https://synergyfinder.fimm.fi/). A summary of synergy scores per cell line is included in the lower right panel. **(C)** Immunoblots showing relative levels of activated ERK (ppERK) in response to vehicle, single agent and combination treatment (24h) for all cell lines (top panel) with quantification shown in the lower panel. **(D)** Cell viability in response to lifirafenib + encorafenib compared with each single agent against a primary *RAS-*mutant AML sample (MC46, left) and cells derived from a primary *RAS*-mutant AML PDX (7638, right). Data points represent either one (MC46) or three replicate experiments. (E) Quantification of immunoblots for MC46 (left) and 7638 (right) in response to vehicle, single agent or combination treatment (24h). Images of immunoblots are provided in Supplementary Figure S2D.

We next assessed the effect of the combination on pathway activity via measurement of double phosphorylated ERK (ppERK, which we have shown correlates strongly with cell viability^22^) 24 hours post-drug treatment. In line with our cell viability results, ppERK was significantly reduced in all cell lines following combination treatment, as compared with single agent and vehicle-treated cells. Compared with vehicle-treated cells, ppERK levels were reduced 0.36-, 1.7– and 1.6-fold in HL-60, OCI-AML3 and THP-1 lines, respectively (Figure 2C). Assessment of ppERK levels also highlighted the potential for paradoxical activation with the use of single RAF inhibitors, with increased pathway activity observed in 4/5 cell lines in response to encorafenib, and 2/5 cell lines following lifirafenib treatment (Figure 2C).

Next, we assessed the efficacy of the combination against two patient-derived *NRAS*-mutant AML samples *ex vivo* (sample details provided in Supplementary Figure S2C). These represented either primary AML cells (MC46, *NRAS*^G12D^) or cells derived from NSGS mice following the primary *in vivo* expansion of patient material (7638, *NRAS*^G12V^). In both cases, the combination was more effective than either single agent (Figure 2D). In line with the decrease in viability, we also observed a decrease in ppERK levels in response to combination treatment compared with single agent and vehicle controls (Figure 2E, Supplementary Figure S2D).

In order to benchmark the efficacy of this combination against direct RAS inhibitors, all five cell lines were also treated with BI-2852 (KRAS switch I/II pocket inhibitor) and BI-3406 (KRAS:SOS1 interaction inhibitor). We observed little to no effect on viability, with BI-3406 failing to reach an IC_50_ in any cell line at 10 μM and BI-2852 achieving an IC_50_ in only 1/5 cell lines (NB4, 7.7 µM) (Supplementary Figure S2E).

### Combining Type I/Type II RAF inhibitors is synergistic against NRAS-mutant AML in vitro

As with the previous combination, we first assessed the single-agent efficacy of Type I inhibitors to guide the selection of combinations to be tested. Only two Type I inhibitors are currently available, SB590885 and GDC-0879. Given the superior single-agent efficacy of SB590885 across all cell lines (Supplementary Figure S3A), this was chosen for combination testing alongside the Type II inhibitor lifirafenib. This combination significantly reduced cell viability compared with single agent treatments in all *NRAS*-mutant cell lines but showed only modest or no improvement in *KRAS*-mutant lines (Figure 3A). This was reflected in the assessment of synergy, with Loewe Additivity scores for *NRAS*-mutant cell lines equating to strong synergy (scores = 27.9, 25.4 and 21.5 for THP-1, HL-60 and OCI-AML3, respectively). Loewe additivity scores for *KRAS*-mutant lines indicated either additivity (NB4, score = 2.4) or antagonism (ML-2, score = –4.9) (Figure 3B).

**Figure 3.**
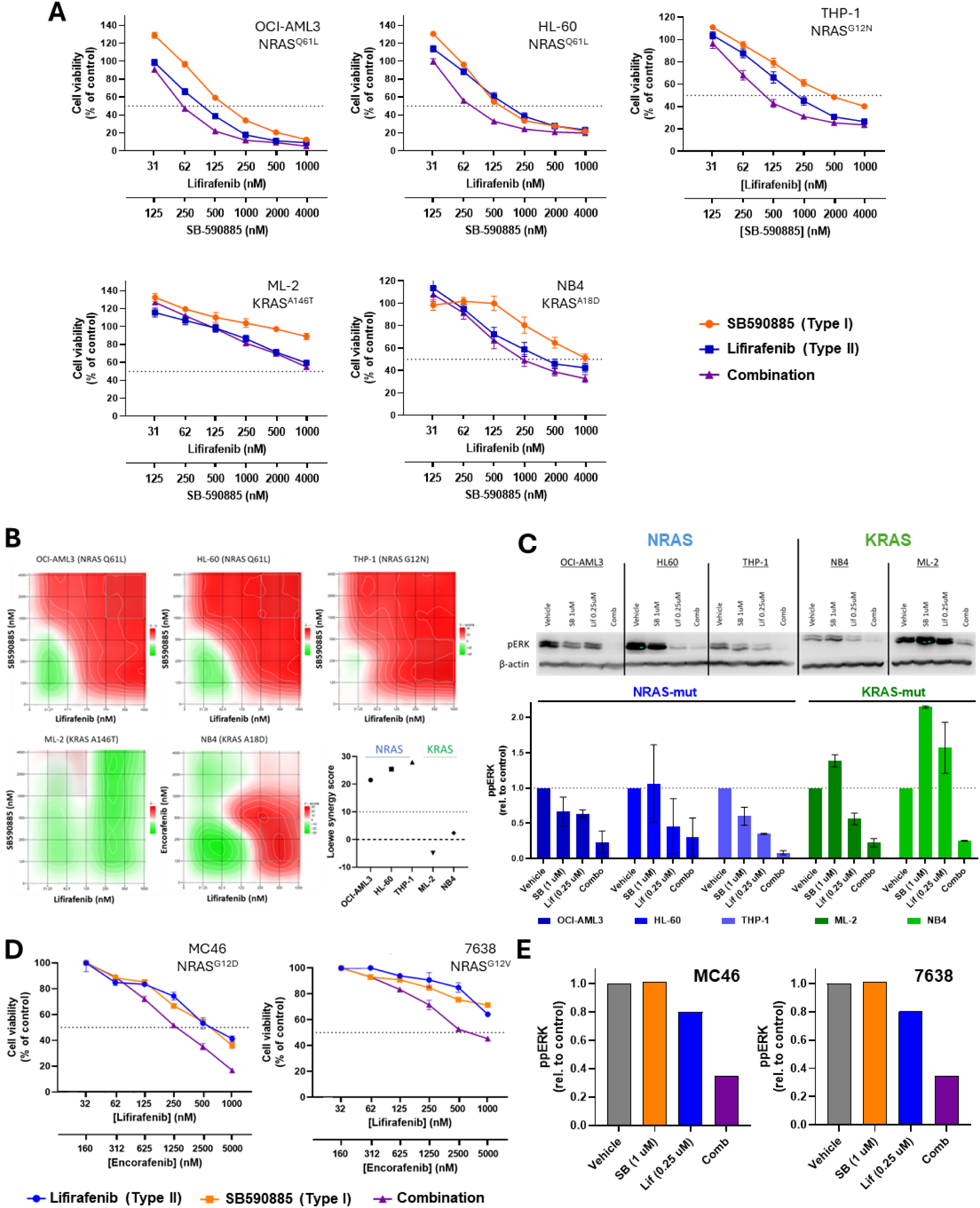
Combining Type I + Type II RAF inhibitors is synergistic against *NRAS*-mutant AML cell lines and patient samples *in vitro*. **(A)** Cell viability in response to lifirafenib (Type II) + SB590885 (Type I) compared with each single agent against five *RAS-*mutant AML cell lines. Cell viability was determined using AlamarBlue at 72 hours post-treatment. Data points represent mean ± SEM from 3 replicate experiments. **(B)** Synergy plots for lifirafenib + SB590885 in all five cell lines tested. Synergy was determined using the Loewe Additivity method, calculated using the SynergyFinder web application (https://synergyfinder.fimm.fi/). A summary of synergy scores per cell line is included in the lower right panel. **(C)** Immunoblots showing relative levels of activated ERK (ppERK) in response to vehicle, single agent and combination treatment (24h) for all cell lines (top panel) with quantification shown in the lower panel. **(D)** Cell viability in response to lifirafenib + SB590885 compared with each single agent against a primary *RAS-*mutant AML sample (MC46, left) and cells derived from a primary *RAS*-mutant AML PDX (7638, right). Data points represent either one (MC46) or three replicate experiments. (E) Quantification of immunoblots for MC46 (left) and 7638 (right) in response to vehicle, single agent or combination treatment (24h). Images of immunoblots are provided in Supplementary Figure S3B.

As with our previous results, synergy in *NRAS*-mutant cell lines was reflected in suppression of ERK pathway activity. Compared with vehicle-treated cells, combination treatment caused a 30.8-, 18.5-or 13.7-fold reduction in ppERK in THP-1, HL-60, and OCI-AML3 cells, respectively (Figure 3C). Interestingly, the ERK pathway activity was also significantly decreased in KRAS-mutant cell lines despite the lack of synergy observed for the combination in cell viability assays.

In line with the combination efficacy observed in *NRAS*-mutant cell lines, this combination was also more effective than single-agent treatment against the two patient-derived samples MC46 and 7638 (Figure 3D). Consistent with the *ex vivo* efficacy experiments, we again observed considerable reduction in ppERK levels in the combination-treated cells compared to single-agent and vehicle treatments (Figure 3D, Supplementary Figure S3B).

### Combined Type I and Type II RAF inhibitors show site-specific efficacy an aggressive RAS-mutant AML in vivo

Using our integrated model, we first calculated the time courses of ppERK levels in AML cells following treatment with Type I and Type II RAF inhibitors, administered either separately or in combination. This analysis was performed for peripheral blood (PB), spleen, and bone marrow (BM) (Figure 4A shows time course for spleen, see Supplementary Figure 4A for PB and BM). We also computed the corresponding areas under the time course curves for each condition (Figure 4A, right panels). The results suggest that this combination of two RAF inhibitors substantially suppressed ERK signaling in PB, spleen and BM, despite strong paradoxical ERK activation observed when the SB590885 RAF inhibitor was administered alone (Fig. 4A).

**Figure 4.**
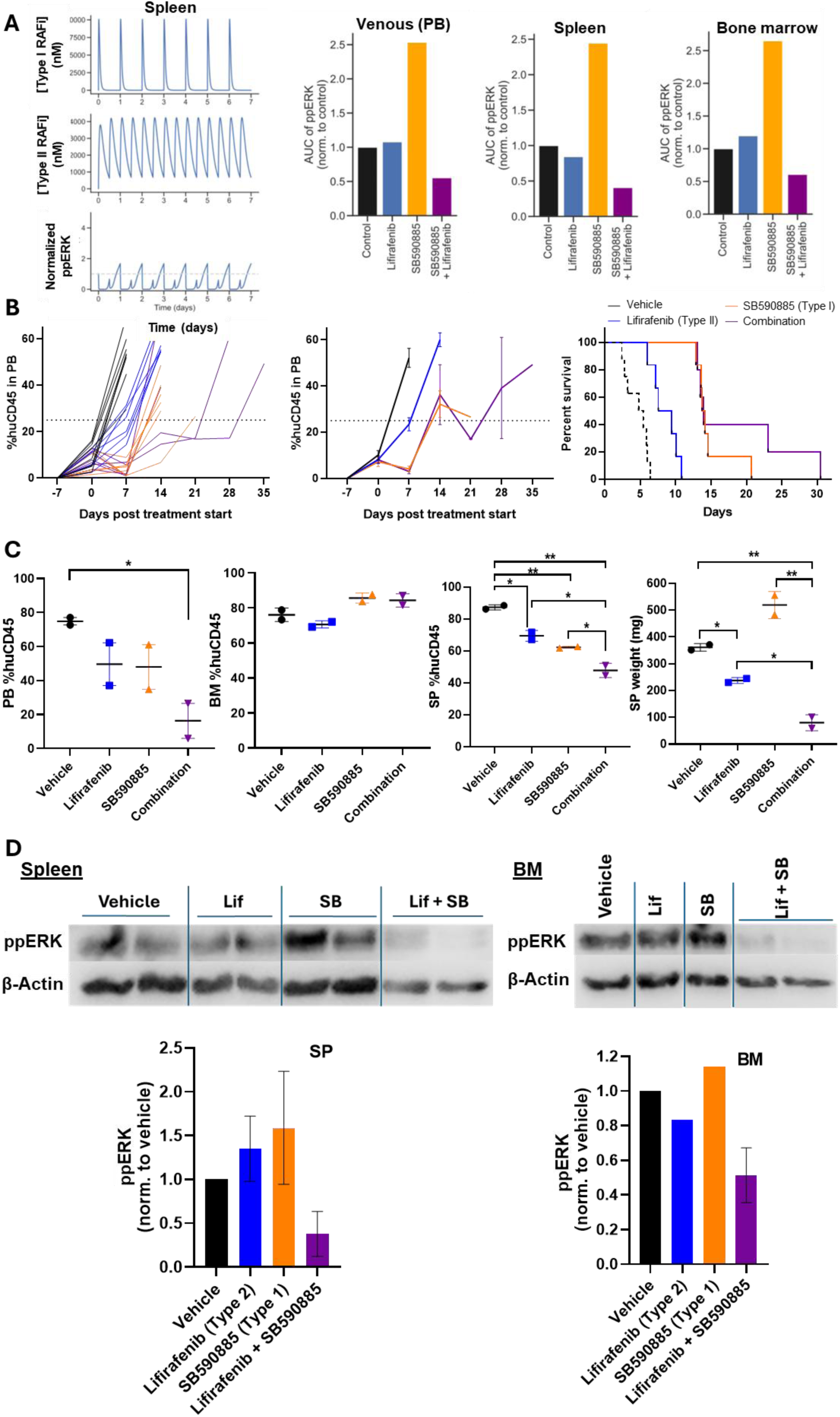
Combining Type I + Type II RAF inhibitors shows site-specific efficacy against an aggressive *NRAS*-mutant AML PDX *in vivo*. **(A)** Predicted phosphorylated ERK (ppERK) levels in response to treatment were determined by our integrated physiologically based pharmacokinetic (PBPK) and structure-based modeling approach. Inhibitor concentrations are given in nanomolar (nM), ppERK levels are normalized to the levels in the untreated condition. The left panel shows time-course predictions of ppERK levels in the spleen over 7 days (see Supplementary Figure S4A for ppERK level predictions in peripheral blood [PB] and bone marrow [BM]). The right panels display the area under the curve (AUC) of ppERK levels for all treatment groups across PB, spleen, and BM. **(B)** Response of AML005 to vehicle, lifirafenib, SB590885 and combination treatment as determined by percent chimerism of mouse versus human CD45^+^ cells in peripheral blood (PB). The percentage of hCD45^+^ cells is shown for individual mice (left) and for median values for each treatment group (middle). Event-free survival (event = 25% hCD45^+^ cells in PB) is shown for all treatment groups (right). **(C)** Data from two representative mice euthanised at Day 10 post-treatment initiation showing leukemia burden by %hCD45^+^ cells in PB, bone marrow (BM, femur + tibia) and spleen (SP), and by spleen weight for all treatment groups. **(D)** Images and quantification of immunoblots of Day 10 spleen (left) and BM (right) samples.

To validate our findings in a clinically relevant preclinical model, we assessed the *in vivo* efficacy of lifirafenib + SB590885 against AML005, an aggressive AML PDX harboring a *KMT2A::MLLT4* fusion with an activating *NRAS*^G12A^ mutation. Doses and schedules for both drugs were chosen based on previously published data in immune-deficient mouse strains.^35,36^ Additionally, we generated an *in silico* physiologically based pharmacokinetic (PBPK) model, taking into account relative drug concentrations in different anatomical sites. Data from the PBPK model was then integrated with our existing structure-based model of MAPK signaling (see Supplementary Methods).^31^ This approach allowed prediction of ppERK levels in each site in response to treatment. Modelling predicted that combination treatment would effectively suppress ppERK in BM and spleen, as well as circulating blasts (Figure 4A, Supplementary Figure S4A).

Given the reported instability of *RAS* mutation frequencies, the presence of the *NRAS* mutation was confirmed between primary and secondary *in vivo* passages (Supplementary Figure S4B). In line with the aggressiveness of this PDX, all cohorts exhibited high leukemia burden in peripheral blood (PB) at commencement of treatment (group median range 7.2-11.1% huCD45^+^ cells in PB).

In contrast to *in vitro* data, SB590885 was more effective than lifirafenib as a single agent and showed similar efficacy to the combination treatment. Disease regression, as measured by %huCD45+ cells in PB, was observed in 5/8 mice for both the SB590885 and combination groups (Figure 4B). Of the evaluable six SB590885 mice remaining after Day 10, five reached event by the end of the second week of treatment (median Leukemia Growth Delay; LGD = 8.5 days), with one animal achieving an LGD of 15 days. While the lifirafenib + SB590885 combination proved tolerable in earlier tolerability testing performed in naïve mice (Supplementary Figure S4C), we did observe adverse treatment effects in 2/6 mice, prompting removal of one animal from the study. Of the five remaining evaluable mice (post Day 10), three reached event by the end of the second week of treatment (median LGD = 8.4 days), with the remaining two animals achieving an LGD of 18 days and 25.4 days (Figure 4B). Median LGD values for all treatment groups are provided in Supplementary Figure S4D.

To assess therapeutic enhancement, or *in vivo* synergy, we applied a statistical threshold of P=0.05 in comparing median EFS for the combination group versus single treatment cohorts. Both single-agent SB590885 and the combination significantly improved survival compared with lifirafenib (P=0.004 and P=0.01, respectively). Despite the increased LGD observed for two mice in the combination group, there was no significant difference in survival compared with SB590885 alone (P=0.28) and therefore no therapeutic enhancement.

Two representative mice from each cohort were euthanised at Day 10 post-treatment initiation to assess leukaemia burden in the spleen and bone marrow (BM). While SB590885 alone and SB590885 + lifirafenib showed similar clearance of circulating blasts in PB, we observed a marked difference between the two groups in their effect on spleen-resident PDX cells at Day 10. All treatments exhibited significantly lower percentages of huCD45+ cells in the spleen, with the combination group having significantly lower percentage of blasts compared with SB590885 alone (P=0.045, Figure 4C). Notably, the spleens from combination mice were significantly smaller (median = 80mg, 0.22-fold reduction vs. vehicle) than all other groups, particularly the SB590885-treated mice which had considerably enlarged spleens (median = 519 mg, 1.44-fold increase vs. vehicle) (Figure 4C). These results suggest that while both SB590885 and the SB590885 + lifirafenib combination effectively clear circulating blasts from the PB, the combination potently targets leukemia residing in the spleen. We next performed immunoblotting on Day 10 spleen and BM samples to determine the effect of each treatment on RAS/RAF/MEK/ERK signaling, measured by ERK phosphorylation (ppERK), in different anatomical niches. In line with spleen weights, we observed a 0.37-fold reduction in ppERK levels in spleen-derived cells from combination-treated mice compared with vehicle controls (Figure 4D). We also observed paradoxical activation of ppERK in the spleens of mice treated with either single agent, which was more pronounced in SB590885-treated samples (Figure 4D).

Despite the efficacy observed in spleen samples, the combination proved ineffective at clearing leukemia from the BM. No significant differences were observed in BM %huCD45+ cells for any treatment group compared with the vehicle-treated cohort (Figure 4B). Interestingly, we observed a 0.51-fold reduction in ppERK levels in the BM of combination-treated mice compared with vehicle controls (Figure 4D).

### Combining Type ***I½*** and Type II RAF inhibitors improves survival of an aggressive RAS-mutant AML in vivo

As in the previous section, we first applied our integrated model to calculate the time courses of ppERK levels in AML cells, this time following treatment with Type I½ and Type II RAF inhibitors, administered either individually or in combination. The analysis was conducted in PB, spleen, and BM (Figure 5A, left panels). As above, we also calculated the corresponding areas under the time course curves for each condition (Figure 5A, right panels). The results suggest that a combination of Type I½ and Type II RAF inhibitors substantially inhibited ERK signaling in PB, spleen and BM.

**Figure 5.**
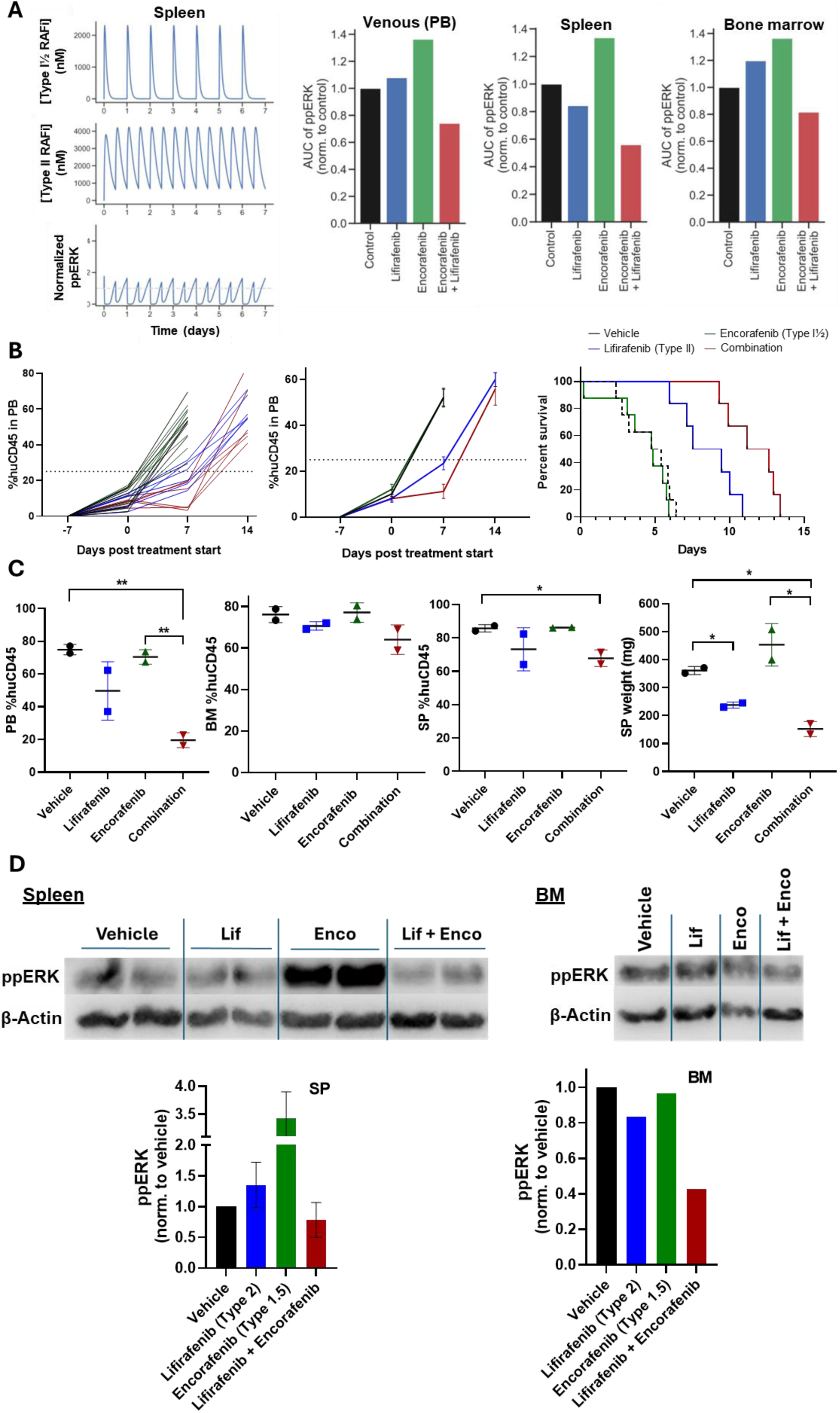
Combining Type I½ + Type II RAF inhibitors synergistically improves survival of an aggressive *NRAS*-mutant AML PDX *in vivo*. (A) Predicted levels of phosphorylated ERK (ppERK) in response to treatment were determined by our integrated physiologically based pharmacokinetic (PBPK) and structure-based modelling approach. Inhibitor concentrations are expressed in nanomolar (nM), ppERK levels are normalized to the level in the untreated condition. The left panel shows time-course predictions of ppERK levels in the spleen over 7 days (see Supplementary Figure S5A for ppERK level predictions in peripheral blood [PB] and bone marrow [BM]). The right panels display the area under the curve (AUC) of ppERK levels for all treatment groups across PB, spleen, and BM. **(B)** Response of AML005 to vehicle, lifirafenib, encorafenib and combination treatment as determined by percent chimerism of mouse versus human CD45^+^ cells in peripheral blood (PB). The percentage of hCD45^+^ cells is shown for individual mice (left) and for median values for each treatment group (middle). Event-free survival (event = 25% hCD45^+^ cells in PB) is shown for all treatment groups (right). **(C)** Data from two representative mice euthanised at Day 10 post-treatment initiation showing leukemia burden by %hCD45^+^ cells in PB, bone marrow (BM, femur + tibia) and spleen (SP), and by spleen weight for all treatment groups. **(D)** Images and quantification of immunoblots of Day 10 spleen (left) and BM (right) samples.

We next assessed the *in vivo* efficacy of lifirafenib in combination with encorafenib in the same PDX model. Inhibitor doses and schedules were again based on previously published data.^37^ As observed for the lifirafenib + SB590885 combination, our integrated PBPK/structure-based model predicted that the combination of lifirafenib + encorafenib would synergistically reduce ppERK levels in AML cells in the BM, spleen and PB (Figure 5A, Supplementary Figure S5A). Results of tolerability testing for this combination are shown in Supplemental Figure S5B.

Despite the high level of engraftment at the beginning of treatment, 2/8 combination-treated mice achieved leukemia regression in the first week of treatment, while regression was not observed in response to either single agent (Figure 5B). The combination achieved a significantly greater LGD (6.83 days) compared with either lifirafenib (4.39 days, P=0.019) or encorafenib (–0.25 days, P<0.001) alone (Supplementary Figure S5C). Using the threshold of P < 0.05, the combination group achieved therapeutic enhancement as the EFS for combination-treated mice was significantly greater than lifirafenib-treated (P=0.015) and encorafenib-treated (P<0.001) cohorts (Figure 5B).

In line with the PB data, encorafenib-treated mice showed equivalent engraftment levels to vehicle-treated mice in both BM and spleen at Day 10. Comparing the combination with lifirafenib alone showed only a modest decrease in the percentage of huCD45+ cells in BM and spleen, despite the combination group exhibiting considerably fewer circulating blasts at this time point (Figure 5C). Interestingly, increased clearance of spleen-resident PDX cells by the combination, as compared with lifirafenib alone, was more evident when assessing raw spleen weight (Figure 5C).

We observed considerable activation of ppERK in the spleen-derived cells of encorafenib-treated mice when compared with the control cohort (Figure 5D), which was in line with the increased spleen weight. Lifirafenib-treated mice showed a slight increase in ppERK levels in the spleen compared with controls, despite the reduction observed in %huCD45+ cells and spleen weight (Figure 5C and 5D). This might be explained by a paradoxical pathway activation by a single RAF inhibitor. Compared with vehicle– and single agent-treated groups, spleen samples from combination-treated mice exhibited a mild reduction in ppERK levels. This was in line with the reduction observed in measurements of blast percentage and weight from spleen samples. Day 10 samples from BM showed a strong reduction (0.42-fold) in ppERK following treatment with the combination compared with vehicle. Surprisingly, we observed a 0.5-fold reduction in ppERK signal for the combination compared with lifirafenib treatment despite there being only a mild difference in %huCD45+ cells in the BM (Figure 5C and D). As we observed a similar decrease in ppERK levels in response to the lifirafenib + SB590885 combination (Figure 4D), it is possible that targeting RAS/RAF/MEK/ERK signaling in the BM is not sufficient to cause cell death due to compensatory factors in the BM niche. Nevertheless, these results suggest that the lifirafenib + encorafenib combination provides marked benefits in therapeutic effect and tolerability when compared with the equivalent lifirafenib + SB590885 treatment.

## Discussion

In this study we showed that dual targeting of RAF by combining discrete, conformation-specific RAF inhibitors is a promising strategy for treating *RAS*-mutant AML. Using our refined, AML-focused model of RAS/RAF/MEK/ERK signaling we predicted that co-inhibition of RAF using either Type I½ + Type II or Type I + Type II inhibitor combinations would synergistically decrease pathway activity. These predictions were first validated in *in vitro* models representing both *NRAS*– and *KRAS*-mutant AML. Importantly, both combination types proved effective against primary *NRAS*-mutant AML samples *ex vivo* and an aggressive pediatric *NRAS*-mutant AML PDX model.

Mutations activating RAS signaling are common in AML, with greater frequencies observed in pediatric patients, and often co-occur with high-risk translocations such as *KMT2A*– and *NUP98*-rearrangements.^38^ The prognostic impact of *RAS-* and RAS pathway mutations in AML remains unclear, depending largely on age, cooperative lesions and mutation type.^13,14,39–41^ Mutations in *KRAS*, *PTPN11* and *NF1* have been reported as independent predictors of poor outcome.^42–46^ Moreover, *RAS* mutations have been increasingly associated with relapse and inferior survival following non-intensive therapies, including hypomethylating agents, FLT3 inhibitors, IDH inhibitors and venetoclax.^5,6,10,11^ Despite the potential benefit of inhibiting RAS pathway activation in AML, previous efforts to target downstream effectors such as MEK have shown only modest efficacy.^47,48^ More recently, inhibition of RAF in *RAS*-mutant AML has been investigated using the pan-RAF inhibitors LY3009120 and belvarafenib, showing modest single-agent activity.^49,50^ To our knowledge, this study is the first to explore dual RAF inhibition in blood cancers, and in doing so we provide a novel, non-obvious and effective therapeutic strategy for treating *RAS*-mutant AML.

A major consideration in the use of inhibitors targeting RAS/RAF/MEK/ERK signaling is the potential for paradoxical activation of downstream signaling, particularly when targeting RAF. This phenomenon results from ligand-induced dimerization and allosteric modulation of the opposing (unbound) monomer to increase kinase activity.^51,52^ Following *in vitro* treatment, paradoxical activation was observed in response to encorafenib and SB590885 when used as single agents. Importantly, in all cases both combination types ablated this effect to synergistically reduce ppERK levels, which is in line with our previous observations in solid tumor models.^22,30^ While the reduction in ppERK levels largely correlated with cell viability in response to both combinations, this was not observed for the lifirafenib + SB590885 combination against *KRAS-*mutant cell lines. This suggests activation of compensatory signaling pathways, however this discrepancy between ppERK levels and cell viability was not observed in response to lifirafinib + encorafenib.

While lifirafenib + SB590885 showed considerable efficacy *in vivo* against AML005, the combination did not significantly improve EFS compared with single-agent SB590885. However, assessment of leukemia burden in the spleens of representative mice at Day 10 showed significantly greater clearance of blasts in response to combination treatment. This was most pronounced in the measurement of spleen weight, where combination treatment decreased weight to approximately the size of non-engrafted immune-deficient mice.^53,54^ Conversely, SB590885 alone increased the spleen weight compared with control mice. While SB590885 treatment appeared to reduce the percentage of hCD45^+^ cells in the spleen compared with controls, the significantly increased spleen size suggests that the number of blasts had increased. This may be explained by the increase in ppERK levels observed in the spleens of SB590885-treated mice and provides important context for the benefit of combining Type I and Type II inhibitors despite observing similar EFS based on peripheral blood values. While our data suggests that Type I + Type II RAFi combinations warrant further investigation, translational relevance of Type I inhibitors is currently limited. To date, no Type I inhibitor has progressed beyond the preclinical stage, and use of these inhibitors as single agents causes strong paradoxical activation of downstream signaling in *RAS*-mutant/*RAF*-WT cancers.^20^

Importantly, the Type I½ and Type II inhibitors used in this study are either under active clinical investigation (lifirafenib) or FDA-approved (encorafenib). This combination showed significantly greater efficacy compared to either single agent, which is striking in the context of a highly aggressive PDX model. As a result of the rapid expansion of AML005 in NSGS mice, the leukemia burden across all cohorts was significantly higher at treatment initiation than the intended ≥ 1% hCD45^+^ cells in PB. Interestingly, despite observing only a modest decrease in %hCD45^+^ cells in BM upon combination treatment, ppERK levels were significantly reduced. This suggests that these agents reach the BM but the effect of RAF inhibition on cell viability is diminished, likely due to microenvironmental factors. One explanation for this may be the supraphysiological levels of stem cell factor (SCF) in NSGS mice.^55^ Binding of SCF to its cognate receptor, c-KIT, can activate signaling pathways such as PI3K-mTOR which can compensate for RAS/RAF/MEK/ERK signaling.^56^ The hypoxic nature of the BM microenvironment may also be a contributing factor. It has been reported that parallels exist between hypoxia and the consequences of RAS activation, with both exhibiting induction of fatty acid scavenging.^57^ This overlap may increase the permissiveness of the BM microenvironment for the growth of *RAS*-mutant AML.

Future investigations into either combination type may benefit from modulation of doses and schedules used *in vivo*. Predictions of the effect of each combination using our integrated PBPK/structure-based model were corroborated by experimental data from mice evaluated at Day 10 during treatment. However, the incorporation of experimental data into our model would refine predictions to inform more precise dosing schedules.

Together, our data suggests that dual targeting of RAF using conformation-specific inhibitors is an effective, well-tolerated strategy for treating *RAS-*mutant AML. These results provide a platform for refining RAFi combinations for clinical use, and a proof-of-concept that may be applied to other hematological malignancies with activated RAS signaling.

## Acknowledgements and Funding

The authors thank the Irish Cancer Society, which fully supported LJ throughout this project through a Translational Research Fellowship (CRF20JON). The authors would also like to thank the National Children’s Research Centre for their significant support of this work through Leadership Award A-18-3 that funded start-up research in the Bond laboratory at Systems Biology Ireland. Additionally, the authors thank and acknowledge the many team members at Systems Biology Ireland and the Irish National Children’s Cancer Service at Children’s Health Ireland at Crumlin for their support and helpful suggestions during this project.

TD was co-funded by Children’s Health Ireland and Research Ireland as part of the Precision Oncology Ireland programme (18/SPP/3522). Work in the Bond laboratory is also funded by Research Ireland grant 20/FFP-P/8844. B.K. was supported by NIH grants R01CA244660 and R01HL171773, EU grant no. 101136926 MULTIR. OR was supported by Science Foundation Ireland 22/PATH-S/10875. Proteomic and phosphoproteomic analyses were supported by the Comprehensive Molecular Analytical Platform (CMAP) under the SFI Research Infrastructure Programme, reference 18/RI/5702. CC was funded by Research Ireland through the SFI Centre for Research Training in Genomics Data Science under Grant number 18/CRT/6214.

The authors would like to thank the members of the Patient-Derived Xenograft Core at the Baylor College of Medicine, TX, USA, who kindly provided the AML-005 sample used in this study. We would also like to thank Dr. Ronald Stam and Susan Arentsen-Peters from the Princess Maxima Center for Pediatric Oncology, Utrecht, The Netherlands, for kindly providing the 7638 sample. Additionally, we thank the team at VIVO Biobank (UK). The MC46 sample and associated data used in this study was provided by VIVO Biobank, supported by Cancer Research UK & Blood Cancer UK (Grant no. CRCPSC-Dec21\100003)

