## Supplemental Data for "A structure-based modelling approach identifies effective drug combinations for RAS-mutant acute myeloid leukemia"

**Jones *et al.***

**Contents:**

- Supplemental Methods
- Supplemental Figures S1-S5
- Supplemental References

### Supplemental Methods

#### Protein extraction

Protein lysates were obtained from cell pellets using an in-house lysis buffer (1M Tris pH7.5, 5M NaCl, and 0.5% (v/v) NP40, H2O, filtered) supplemented with protease and phosphatase inhibitors (COmplete Mini Protease Inhibitor Cocktail – Roche 11836153001; PhosSTOP Phosphatase Inhibitor Cocktail – Roche 4906837001).

#### Immunoblotting

15 to 20µg of protein/sample were mixed (4:1 v/v) with a mix of DL-Dithiothreitol (DTT) plus NuPAGE LDS Sample Buffer (1:10), incubated for 10 minutes at 95°C and then resolved on 10 to 12% acrylamide electrophoresis (running buffer: 0.25 M Tris-HCL pH 8.8, 1.9 M Glycine, 0.2% SDS) before being transferred to a PVDF membrane using a wet transfer method (transfer buffer: 200 nM Tris-HCL pH 7.5, 50 mM EDTA, 1M NaCl). The membrane was washed with tris buffered saline tween (TBST) before blocking with 5% (w/v) skimmed milk powder in TBST shaking for 1.5h. The membrane was then incubated overnight with primary antibody at 4°C. After washing with TBST, the membrane was incubated for one hour with a HRP-conjugated secondary antibody to detect the binding between the primary antibody and the protein. The chemoluminescent signal was detected by using homemade ECL and exposure in Advanced Molecular Vision Chemi Image Unit of the ChemoStar Imager (INTAS Science Imaging Instruments GmbH). Polyclonal rabbit anti-human mitogen-activated protein (MAP) kinase [extra-cellular signal-regulated kinase (ERK) 1 & 2] antibody (#M5670) and monoclonal mouse anti-human MAP kinase, activated (diphosphorylated ERK-1 & 2) antibody (#8159) were both purchased from Sigma-Aldrich, Burlington, MA, USA. Monoclonal rabbit anti-human β-actin (#4970) was purchased from Cell Signaling Technology, Danvers, MA, USA.

**Mass Spectrometry**

Prior to Mass Spectrometry analysis, lysates and enriched samples were resuspended in 0.1% formic acid buffer. Samples were run on a Bruker timsTof Pro mass spectrometer connected to an Evosep One liquid chromatography system. Tryptic peptides were resuspended in 0.1% formic acid and each sample was loaded on to an Evosep tip and separated on a Evosep EV1106 Endurance Column – 15 cm x 150 um, 1.9 um. The mass spectrometer was operated in positive ion mode with a capillary voltage of 1500 V, dry gas flow of 3 l/min and a dry temperature of 180 °C. All data was acquired with the instrument operating in trapped ion mobility spectrometry data dependent acquisition mode (TIMS DDA) mode. Trapped ions were selected for ms/ms using parallel accumulation serial fragmentation (PASEF). A scan range of (100-1700 m/z) was performed at a rate of 5 PASEF MS/MS frames to 1 MS scan with a cycle time of 1.03s 8. Chromatography Buffers were as follows: Buffer B: 99.9% acetonitrile, 0.1% formic acid. Buffer A: 99.9% water, 0.1% formic acid. All solvents are LCMS grade.

**Proteomic/phospho-proteomic sample preparation**

Samples first underwent trypsin digestion; DTT was added to 300 µg of cell lysate to reach a final concentration of 100mM and then placed on an Eppendorf® Thermomixer® R (model T3317) for shaking (750 rpm) at 30°C for 30 minutes. Iodoacetamide was added to the sample to reach a final concentration of 20 mM then incubated for 30 minutes in the thermomixer (750 rpm, 30°C). Samples were then diluted 1 in 4 (v/v) using 50 mM Tris-HCl and trypsin (Promega, V5111) was added before overnight shaking (750 rpm), at 37°C. Samples were then collected and formic acid added (1% final volume), followed by a brief vortex. C18 tips (Pierce™ C18 Spin Tips & Columns, cat number: 87784) were activated using 100 µL of 80% acetonitrile (ACN) and 0.1% trifluoracetic acid (TFA) buffer before columns were spun at 3,000 rpm for 1 minute The tips where then equilibrated by adding 100 µL of 0.1% trifluoracetic acid (TFA) and centrifuged at 3000 rpm for 1 minute.  600 µL of sample was loaded to the column and centrifuged for 2 minutes at 850g (in 3 different rounds, i.e., 200 µL + centrifugation, then repeat). The column was then washed with a 0.1% TFA buffer before spinning the column at 850g for 2 minutes. The sample was eluted in a fresh tube using a buffer containing 0.1% TFA and 80% ACN and centrifugation for 2 minutes at 850g. The post-cleaning eluate was used for phospho-enrichment.

**Mass Spectrometry Data Analysis**

For analysis of MS output data, peptide mapping was initially performed with MaxQuant (release 2.0.1.0) using the Homo sapiens subset of the Uniprot Swissprot database with specific parameters for TIMS data dependent acquisition (TIMS-DDA). The MaxQuant output file was imported into the Perseus (version 1.6.15.0) environment for protein quantification. LFQ (Label Free quantitation) intensities were loaded as main columns. Reverse proteins and proteins only identified by site were filtered out from further analysis. LFQ values were then log2 transformed. The data frame was then split into smaller datasets by group of samples (e.g., condition v control) allowing filtering of proteins that were not identified in more than one replicate. Missing values were imputed based on the normal distribution with a width of 0.3 and a down shift of 1.8. These condition-specific (such as control condition and treated condition) data frames were then exported and used for further analysis.

For phosphoproteome analysis, label-free data was loaded into Perseus using ‘Intensity X_1’, ‘Intensity X_2’, ‘Intensity X_3’ (i.e., data for the same sample and site for peptides that are mono-, di-, or tri-phosphorylated) as main columns. In addition to reverse proteins and proteins only identified by site, potential contaminants, and proteins with localization probability < 0.75 were excluded from further analysis. The built-in function ‘Expand site table’ was then used to rearrange the columns to permit analysis of differing levels of phosphorylation (i.e., single, double, triple-phosphorylated). Following log2 transformation, the position within the protein, known sites of phosphorylation, and linear motifs were all added to the table based on PhosphoSitePlus data (https://www.phosphosite.org). Similar to the whole proteome data analysis, the data frame was divided based on the condition, proteins filtered based on the number of replicates in which they appeared before imputing the missing values and exporting the data.

**Kinase-Substrate enrichment analysis (KSEA)**

KSEA analysis (version 0.99.0) was performed in Jupyter Notebook coupled with R version 4.4.1. As previously described[1], this approach allows inference of kinase activity based on the collective phosphorylation changes of their identified substrates, as determined from previously analyzed curated data sets. Files that were previously processed in Perseus and R (as described above) were modified to fit the KSEA pipeline requirements, as follows: "Protein" the Uniprot ID for the parent protein, "Gene" the HUGO gene name for the parent protein, "Peptide" the peptide sequence, “Residue.Both" all phospho-sites from that peptide, separated by semicolons (if applicable, these were formatted as the single amino acid abbreviation with the residue position (e.g., S102)), "p" the p-value of that peptide, and "FC" the fold change (not log-transformed). Arguments necessary to run the pipeline are as follow: ‘NetworKIN’ (TRUE or FALSE) to define if NetworKIN prediction should be included or not, ‘m.cutoff’ (numeric value from 1 to infinity) that indicates the minimum number of substrates to be included in the barplot output, and ‘p.cutoff’ (numeric value between 0 and 1) that indicates the p-value cut-off for inclusion of significant kinases in the bar plot.

**Cell viability assays**

For assessment of drug responses, cells were seeded in 96-well U-bottom plates (10,000 cells/well for cell lines and 50,000 cells/well for primary and PDX samples) prior to addition of inhibitors. Cell viability was assessed at 72h using the Resazurin assay (made in house; 10X stock: resazurin 75 mg, methylene blue 12.5 mg, potassium hexacyanoferrate (III) 164.5 mg, potassium hexacyanoferrate 211 mg in 50 mL PBS). Cells were incubated with Resazurin solution (1:10 v/v) for 4h and fluorescence measured at 560/590 nm using a SpectraMax M3 plate reader (Molecular Devices) SoftMaxPro 6.2.2.

Combination experiments were performed using both fixed-ratio and matrix (6x6) treatments. The Loewe Additivity method was used to determine combination effect, given the shared intracellular target of both agents. Loewe Additivity scores were calculated using the SynergyFinder web application (v3.0; https://synergyfinder.fimm.fi/).

**RAS mutation detection**

Polymerase chain reaction (PCR) products for exon 2 of the *NRAS* gene was amplified from genomic DNA (gDNA) using Phusion High-Fidelity DNA Polymerase (New England BioLabs, Hitchin, UK) according to manufacturer's instructions, with 35 cycles of amplification and an annealing temperature of 60.5°C. Primer sequences used were FWD: 5'-GCTCGCCAATTAACCCTGATTAC-3', REV: 5'-TGGGTAAAGATGATCCGACAAGTGA-3'. Sanger sequencing was performed by Eurofins Genomics (Ebersburg, Germany).

**Physiologically Based Pharmacokinetic Model (PBPK)**

We consider the following 16 compartments in the model: depot for oral administration, depot for intraperitoneal injection, lung, arterial, venous, adipose, muscle, liver, gut, spleen, heart, brain, kidney, skin, bone marrow, and the rest of the body. Total amounts of the drug (in milligrams) in each compartment are denoted as $D_{PO}$, $D_{IP}$, $A_{lu}$, $A_{ar}$, $A_{ve}$, $A_{ad}$, $A_{mu}$, $A_{li}$, $A_{gu}$, $A_{sp}$, $A_{he}$, $A_{br}$, $A_{ki}$, $A_{sk}$, $A_{BM}$, $A_{re}$. Concentration $C_i$ of the drug in compartment $i$ is calculated as $C_i = A_i/V_i$, where $V_i$ is volume of this compartment. The blood flows through lungs, adipose, muscle, gut spleen, heart, brain, kidney, skin, bone marrow and rest of the body are denoted as, $Q_{lu}$, $Q_{ad}$, $Q_{mu}$, $Q_{gu}$, $Q_{sp}$, $Q_{he}$, $Q_{br}$, $Q_{ki}$, $Q_{sk}$, $Q_{BM}$, $Q_{re}$, respectively. For the liver, the flow through hepatic artery is denoted as $Q_{hepa}$, and hepatic venous flow is $Q_{li} = Q_{hepa} + Q_{gu} + Q_{sp}$. Coefficient $BP$ reflects blood to plasma ratio, coefficient $f_{up}$ reflects fraction of the drug unbound in plasma, coefficients $K_p$ reflect tissue to plasma partition coefficients for each compartment.

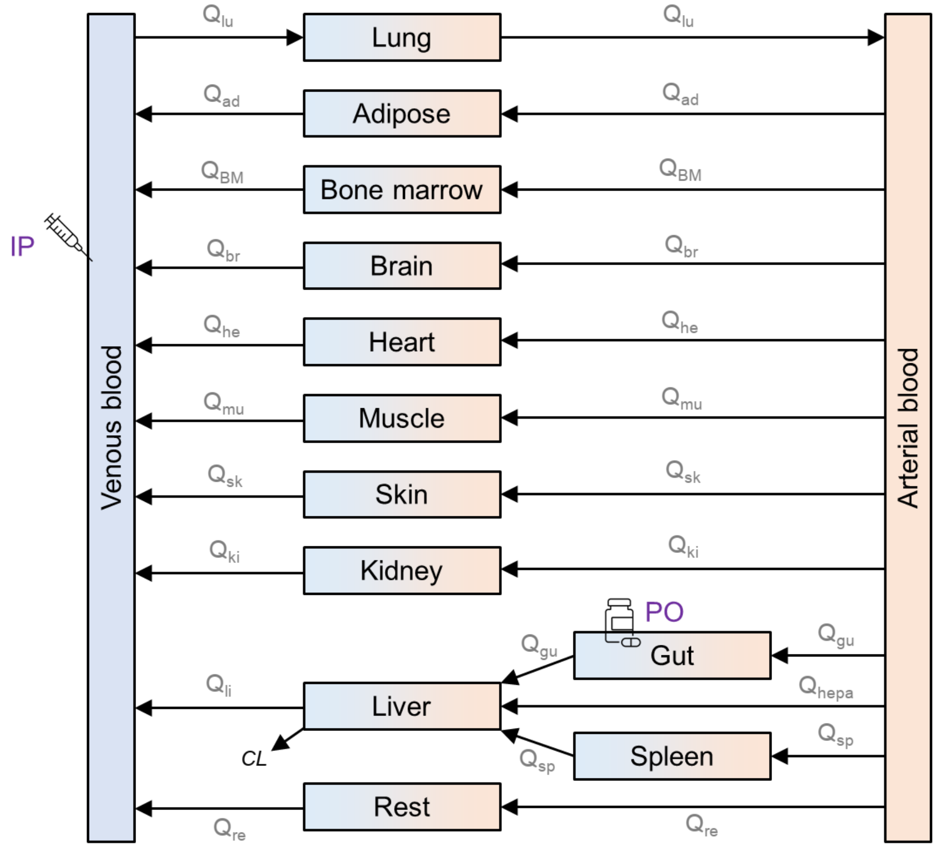

The drug from oral administration (PO) comes to the gut, the drug from intraperitoneal injection (IP) comes to venous blood. We neglect renal clearance of drugs, and take into account only clearance in liver, denoted as $CL$. In such case, PBPK equations read as follows:

| | |
|---|---|
| $\frac{dD_{PO}}{dt} = -K_{aPO} \cdot D_{PO}$ | (1) |
| $\frac{dD_{IP}}{dt} = -K_{aIP} \cdot D_{IP}$ | (2) |
| $\frac{dA_{lu}}{dt} = Q_{lu} \cdot \left( C_{ve} - C_{lu} \cdot \frac{BP}{K_{plu}} \right)$ | (3) |
| $\frac{dA_{ar}}{dt} = Q_{lu} \cdot C_{lu} \cdot \frac{BP}{K_{plu}} - \left( Q_{ad} + Q_{BM} + Q_{br} + Q_{he} + Q_{mu} + Q_{sk} + Q_{ki} + Q_{hepa} + Q_{gu} + Q_{sp} + Q_{re} \right) \cdot C_{ar}$ | (4) |

| | |
|---|---|
| $$\frac{dA_{ve}}{dt} = K_{aIP} \cdot D_{IP} + Q_{ad} \cdot C_{ad} \frac{BP}{K_{pad}} + Q_{mu} \cdot C_{mu} \frac{BP}{K_{pmu}} + Q_{li} \cdot C_{li} \frac{BP}{K_{pli}} + Q_{he} \cdot C_{he} \frac{BP}{K_{phe}} + Q_{br} \cdot C_{br} \frac{BP}{K_{pbr}} + Q_{ki} \cdot C_{ki} \frac{BP}{K_{pki}} + Q_{sk} \cdot C_{sk} \frac{BP}{K_{psk}} + C_{BM} \frac{BP}{K_{pBM}} + Q_{re} \cdot C_{re} \frac{BP}{K_{pre}} - Q_{lu} \cdot C_{ve}$$ | (5) |
| $$\frac{dA_{ad}}{dt} = Q_{ad} \cdot \left(C_{ar} - C_{ad} \cdot \frac{BP}{K_{pad}}\right)$$ | (6) |
| $$\frac{dA_{mu}}{dt} = Q_{mu} \cdot \left(C_{ar} - C_{mu} \cdot \frac{BP}{K_{pmu}}\right)$$ | (7) |
| $$\frac{dA_{li}}{dt} = Q_{hepa} \cdot C_{ar} + Q_{gu} \cdot C_{gu} \frac{BP}{K_{pgu}} + Q_{sp} \cdot C_{sp} \frac{BP}{K_{psp}} - Q_{li} \cdot C_{li} \frac{BP}{K_{pli}} - C_{li} \cdot f_{up} \cdot CL$$ | (8) |
| $$\frac{dA_{gu}}{dt} = K_{aPO} \cdot D_{PO} + Q_{gu} \cdot \left(C_{ar} - C_{gu} \cdot \frac{BP}{K_{pgu}}\right)$$ | (9) |
| $$\frac{dA_{sp}}{dt} = Q_{sp} \cdot \left(C_{ar} - C_{sp} \cdot \frac{BP}{K_{psp}}\right)$$ | (10) |
| $$\frac{dA_{he}}{dt} = Q_{he} \cdot \left(C_{ar} - C_{he} \cdot \frac{BP}{K_{phe}}\right)$$ | (11) |
| $$\frac{dA_{br}}{dt} = Q_{br} \cdot \left(C_{ar} - C_{br} \cdot \frac{BP}{K_{pbr}}\right)$$ | (12) |
| $$\frac{dA_{ki}}{dt} = Q_{ki} \cdot \left(C_{ar} - C_{ki} \cdot \frac{BP}{K_{pki}}\right)$$ | (13) |
| $$\frac{dA_{sk}}{dt} = Q_{sk} \cdot \left(C_{ar} - C_{sk} \cdot \frac{BP}{K_{psk}}\right)$$ | (14) |
| $$\frac{dA_{BM}}{dt} = Q_{BM} \cdot \left(C_{ar} - C_{BM} \cdot \frac{BP}{K_{pBM}}\right)$$ | (15) |
| $$\frac{dA_{re}}{dt} = Q_{re} \cdot \left(C_{ar} - C_{re} \cdot \frac{BP}{K_{pre}}\right)$$ | (16) |

The table below shows compartment-specific parameters (Kp values were calculated using Rodgers & Rowland method[2]). Some studies suggest that the bone marrow functions as a "pharmacological sanctuary," where drug penetration is limited due to protective mechanisms within the microenvironment.[3] This sanctuary effect is attributed to factors such as the expression of drug-metabolizing enzymes like CYP3A4 by bone marrow stromal cells[4], which can inactivate therapeutic agents, and the presence of drug efflux transporters like P-glycoprotein and MRP1 that reduce intracellular drug accumulation.[5] To reflect this reduced drug penetration in the bone marrow compartment, the Kp values were decreased by a factor of 1.25.

The volumes and blood flows were taken from the literature[6,7]:

| Compartment | Volume, L | Blood flow, L/h | Kp, Lifirafenib | Kp, Encorafrenib |
|---|---|---|---|---|
| Lung | 2.06E-04 | 1.2000 | 1.7710 | 1.7715 |
| Arterial | 7.42E-04 | 1.2000 | 1.0000 | 1.0000 |
| Venous | 1.47E-03 | 1.2000 | 1.0000 | 1.0000 |
| Adipose | 2.49E-03 | 0.0134 | 6.8305 | 4.4789 |
| Muscle | 1.20E-02 | 0.1604 | 0.7998 | 0.8541 |
| Liver ($Q_{hepa}$ for flow) | 2.17E-03 | 0.0745 | 1.3660 | 1.3904 |
| Gut | 1.27E-03 | 0.2642 | 2.6753 | 2.6352 |
| Spleen | 1.31E-04 | 0.0159 | 0.7682 | 0.8202 |
| Heart | 1.42E-04 | 0.0494 | 1.2038 | 1.2361 |
| Brain | 5.42E-04 | 0.0803 | 2.4274 | 2.4098 |
| Kidney | 5.07E-04 | 0.2253 | 1.2931 | 1.3201 |
| Skin | 4.58E-03 | 0.1067 | 3.9718 | 3.8513 |
| Bone Marrow | 1.77E-03 | 0.1523 | 0.4800 | 0.5600 |
| Rest | 1.98E-03 | 0.0577 | 1.0000 | 1.0000 |
| Total | 3.00E-02 | 1.2000 | | |

According to conservation laws the following equality must be satisfied:

$$Q_{lu} = Q_{ad} + Q_{BM} + Q_{br} + Q_{he} + Q_{mu} + Q_{sk} + Q_{ki} + Q_{hepa} + Q_{gu} + Q_{sp} + Q_{re} \quad (17)$$

To estimate intrinsic hepatic clearance $CL$, which enters equation (8), we used the well-stirred model [doi:10.1124/dmd.106.013359]. This model connects apparent hepatic clearance $CL_H$, which is usually reported in literature, with intrinsic hepatic clearance $CL$

$$CL_H = Q_{li} \cdot \left( \frac{CL \cdot f_{up}}{Q_{li} + CL \cdot f_{up}} \right) \quad (18)$$

Apparent hepatic clearance $CL_H$ was rescaled from human data using allometric scaling $CL_H \propto W^{0.75}$, where $W$ is weight. These values can be adjusted based on obtained PK curves.

Using equation (18), allometric scaling and publicly available data[8], we estimated other compound-specific parameters as follows:

| Parameter | Value for Encorafenib | Value for Lifirafenib | Value for SB590885 |
|---|---|---|---|
| Clearance, $CL$, L/h | 0.9312 | 3.1093 | Same as Encorafenib |
| Blood-to-plasma, BP | 0.5800 | 0.6000 | Same as Encorafenib |
| $K_{aPO}$, 1/h | 0.7200 | 0.4800 | - |
| $K_{aIP}$, 1/h | - | - | 1.4400 |
| $f_{up}$ | 0.1400 | 0.0050 | Same as Encorafenib |

$K_{aPO}$ were estimated based on $T_{max}$ times as $K_{aPO} = 1.44/T_{max}$. We did not rescale $K_a$ from human to mice using allometric scaling, since (i) absorption rates of other RAF inhibitors lie in the range of 0.5-1 L/h, not 5 L/h, and (ii) absorption rates are not well scaled with weight, unlike clearance rates. $K_{aPO}$ for SB590885 was taken twice higher than $K_{aPO}$ for encorafenib.

All unknown parameters for SB590885 are suggested to take from encorafenib. Initial values for $D_{PO}$ and $D_{IP}$ are taken from dosing regime.

**Integration of the PBPK and structure-based MAPK signaling models**

To predict ERK phosphorylation dynamics in the venous blood, spleen, and bone marrow compartments, we integrated a physiologically based pharmacokinetic (PBPK) model with our structure-based MAPK signaling model. This model is adapted from our previous work[9], with the PI3K-AKT module omitted for simplicity. The resulting model captures key intracellular processes, including ERBB receptor dimerization, recruitment of adaptor proteins to the plasma membrane, RAS activation via SOS1-mediated GDP-to-GTP exchange, RAF dimerization, ERK activation, and negative feedback loops from phosphorylated ERK to RAF, SOS1, and ERBB receptors. Allosteric effects induced by conformation-specific RAF inhibitors (RAFi) are quantitatively represented using thermodynamic factors.[10]

In the previous structure-based model[9], RAFi concentrations were fixed over time. Here, we dynamically updated RAFi concentrations by directly linking the RAFi concentration variables to the time-resolved drug concentration profiles predicted by the PBPK model in the relevant compartments. This allowed the integrated model to capture the temporal dynamics of the drug exposure.

To match the experimental dosing regimen in mice, SB590885 (Type I RAFi) was administered intraperitoneally every 24 hours, encorafenib (Type 1½ RAFi) was administered orally every 24 hours, and lifirafenib (Type II RAFi) was administered orally every 12 hours.

The integrated model was built using the rule-based PySB framework.[11] The SBML files of the model used in this study are available in Supplementary Information.

### Supplemental Figures

#### Figure S1

**A**

| Cell line | RAS mutation | Additional alterations | Age (Yrs) | Sex |
|---|---|---|---|---|
| OCI-AML3 | *NRAS*$^{Q61L}$ | *DNMT3A*$^{R882C}$<br>NPM1c+ | 57 | M |
| HL-60 | *NRAS*$^{Q61L}$ | *TP53*$^{null}$<br>*CDKN2A* Hom c.238C>T | 36 | F |
| THP-1 | *NRAS*$^{G12N}$ | *KMT2A::MLLT3*<br>*CSNK2A1::DDX39B*<br>*TP53* Het c.520_545del26 | 1 | M |
| NB4 | *KRAS*$^{A18D}$ | *PML::RARA*<br>*TP53* Hom c.743G>A | 23 | F |
| ML-2 | *KRAS*$^{A146T}$ | *KMT2A::AFDN* | 26 | M |

**B**

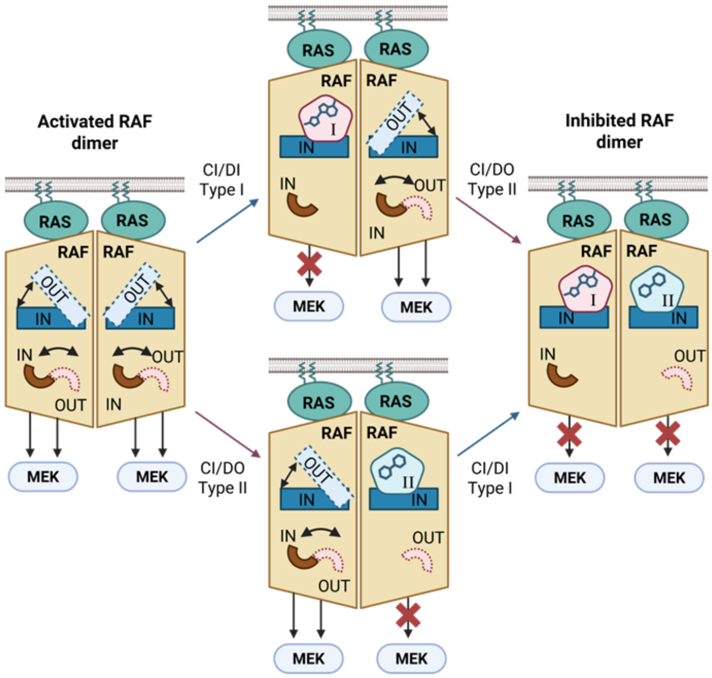

**Figure S1. (A)** Characteristics of the AML cell lines used in this study. **(B)** Cartoon depicting the binding preferences and effects on downstream signaling upon treatment with Type I, Type II and Type I + Type II RAF inhibitors.

Figure S2

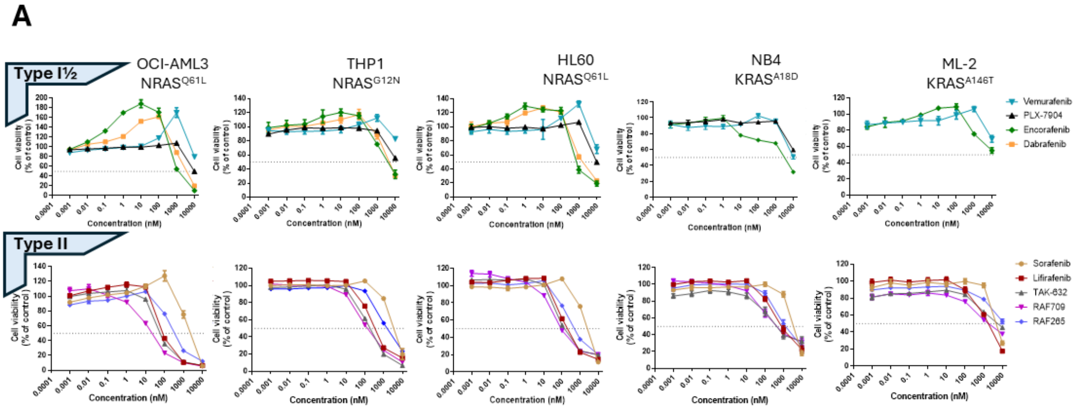

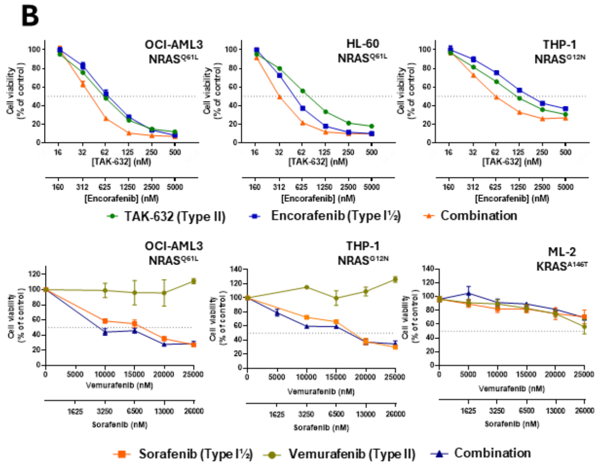

C

| Sample | RAS mutation | Other alterations | Sex | Age @ Dx (Yrs) |
|---|---|---|---|---|
| 7638 | NRAS G12V | KMT2A::KIAA1524 | F | 0.4 |
| MC46 | NRAS G12D | Inv(16)(p13q22) | M | 16 |

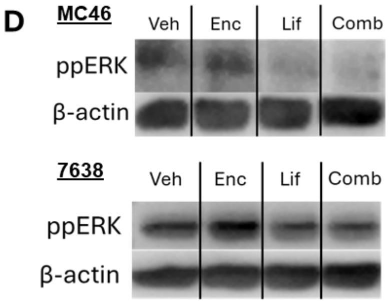

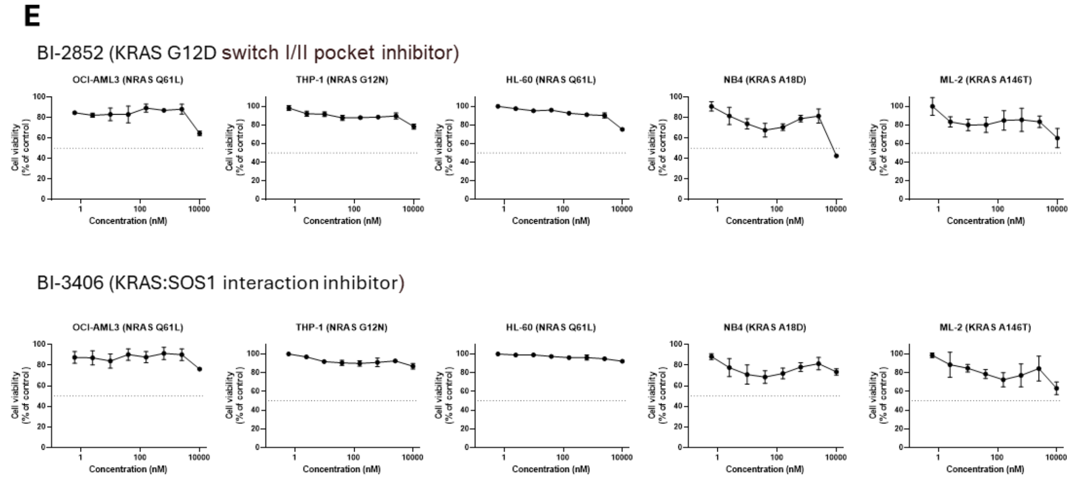

**Figure S2. (A)** Single-agent dose-response curves for all Type I½ and Type II RAF inhibitors tested against five *RAS*-mutant cell lines. **(B)** Cell viability in response to TAK-632 (Type II) + encorafenib (Type I½) compared with each single agent against representative *RAS*-mutant cell lines. **(C)** Characteristics of primary AML samples used for *ex vivo* drug treatments. **(D)** Immunoblots for MC46 (top) and 7638 (bottom) measuring ppERK levels to vehicle, single agent or combination treatment (24h). **(E)** Single-agent dose response for RAS-specific inhibitors. For cell viability experiments, cell viability was determined using AlamarBlue at 72 hours post-treatment. Data points represent mean + SEM from 3 replicate experiments.

Figure S3

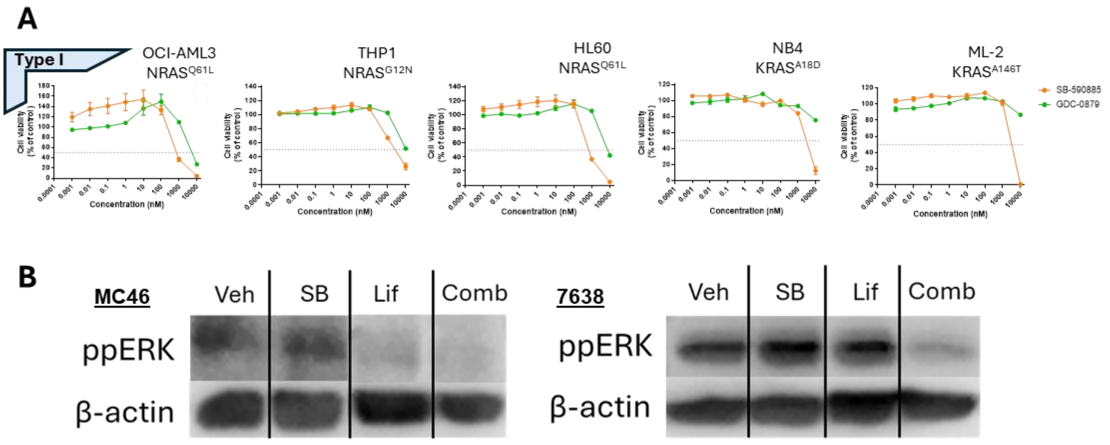

**Figure S3. (A)** Single-agent dose-response curves for all Type I RAF inhibitors tested against five *RAS*-mutant cell lines. **(B)** Immunoblots for MC46 (left) and 7638 (right) measuring ppERK levels to vehicle, single agent or combination treatment (24h).

Figure S4

A

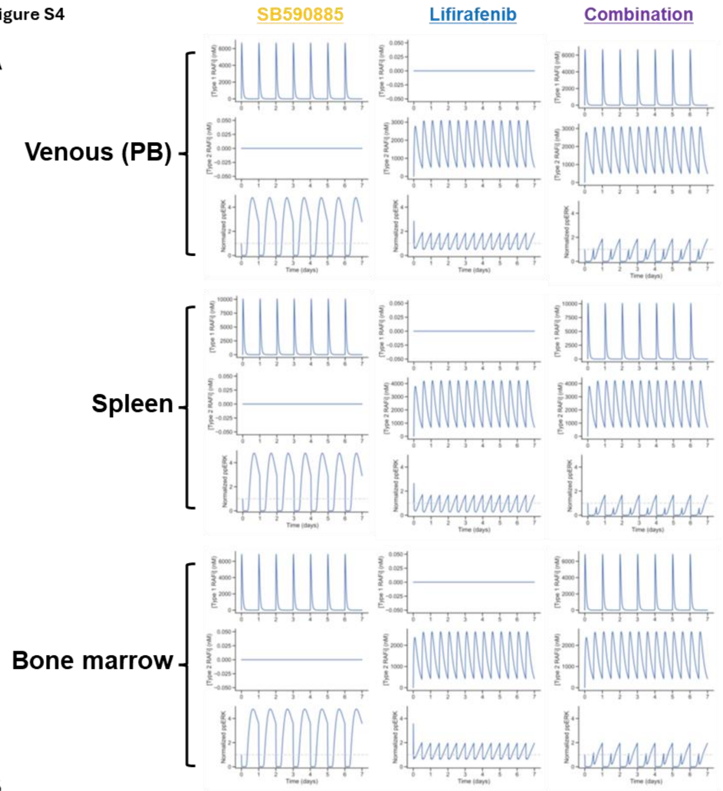

B

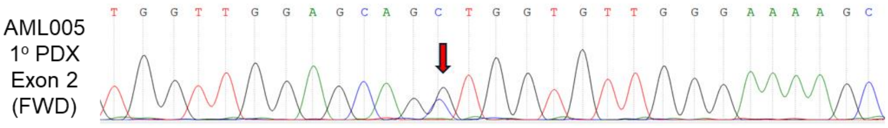

C

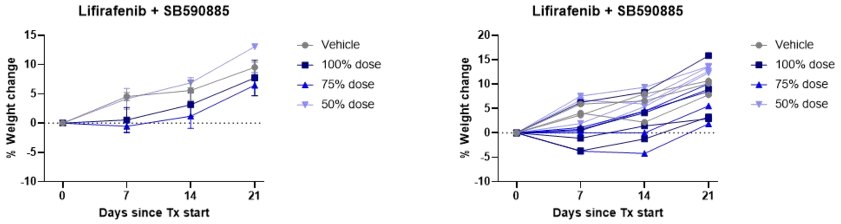

D

| Treatment group | Median event-free survival (**EFS**) (Days) | Median leukemia growth delay (**LGD**) (T-C; Days) |
|---|---|---|
| Vehicle | 5.08 | - |
| Lifirafenib | 8.50 | 4.39 |
| SB590885 | 13.79 | 8.71 |
| Combination | 13.99 | 8.91 |

**Figure S4. (A)** Area under the curve (AUC) for prediction of ppERK levels over time during treatment in venous blood (PB), spleen and bone marrow. **(B)** Sanger sequencing showing point mutation in Exon 2 of *NRAS* in AML005. **(C)** Tolerability data for lifirafenib + SB590885 treatment as determined by weight loss over time. **(D)** Leukemia growth delay (LGD) values calculated for AML005 treated with combination or single agents.

Figure S5

**A**

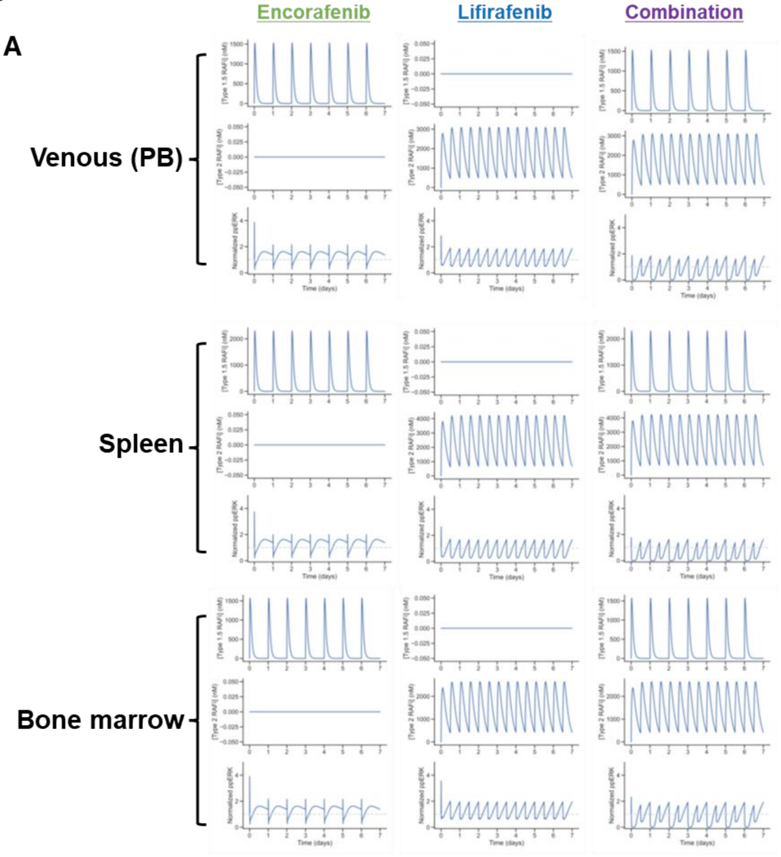

**B**

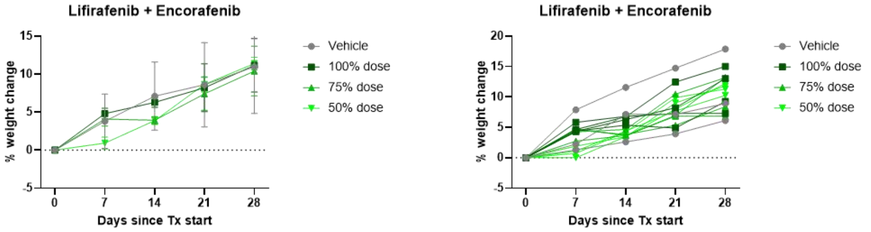

**C**

| Treatment group | Median event-free survival (**EFS**) (Days) | Median leukemia growth delay (**LGD**) (T-C; Days) |
|---|---|---|
| Vehicle | 5.08 | - |
| Lifirafenib | 8.50 | 4.39 |
| Encorafenib | 4.83 | -0.25 |
| Combination | 11.91 | 6.83 |

**Figure S5. (A)** Area under the curve (AUC) for prediction of ppERK levels over time during treatment in venous blood (PB), spleen and bone marrow. **(B)** Tolerability data for lifirafenib + encorafenib treatment as determined by weight loss over time. **(D)** Leukemia growth delay (LGD) values calculated for AML005 treated with combination or single agents.
